# Rapid and sensitive detection of native glycoRNAs

**DOI:** 10.1101/2023.02.26.530106

**Authors:** Helena Hemberger, Peiyuan Chai, Charlotta G. Lebedenko, Reese M. Caldwell, Benson M. George, Ryan A. Flynn

**Author notes:** Corresponding Author: Ryan A. Flynn.

## Abstract

Chemical tools enable precise characterization of many biopolymers, including glycoconjugates. Metabolic chemical reporters enabled the discovery of glycoRNAs, however they have certain limitations due the requirement of having living cells to incorporate the modified sugar. Here we develop a periodate oxidation and aldehyde ligation method to detect and characterize native sialoglycoRNAs, termed rPAL. With optimized RNA biochemistry to enhance recovery and analysis of small RNAs, we show rPAL is at least an order of magnitude more sensitive than previous methods for detecting sialoglycoRNAs. These improvements allow rPAL to detect sialoglycoRNA from human clinical samples as demonstrated by defining the abundance and patterns of sialoglycoRNAs from sorted populations of peripheral blood mononuclear cells. The sensitivity, robustness, and flexibility of rPAL will allow greater access towards characterizing glycoRNA biology.

## Introduction

Metabolic chemical reporters (MCR) have been widely implemented to investigate various aspects of all biopolymers. We recently showed that a sialic acid specific MCR, N-azidoacetylmannosamine-tetraacylated (Ac_4_ManNAz, **Figure 1A**), is able to incorporate into sialylated N-glycans that are conjugated to small noncoding RNAs in mammalian cells^1^. These glycoRNAs, or more precisely sialoglycoRNAs, were produced from a wide variety of cell types, both in vitro and in vivo, and critically were demonstrated to be presented on the external surface of living cells^1^. Most of our previous work critically relied on metabolic incorporation of Ac_4_ManNAz, leveraging its incorporation into sialic aids and the chemical selectively of the azido-group for bioorthogonal labeling with copper-free click reagents.

**Figure 1.**
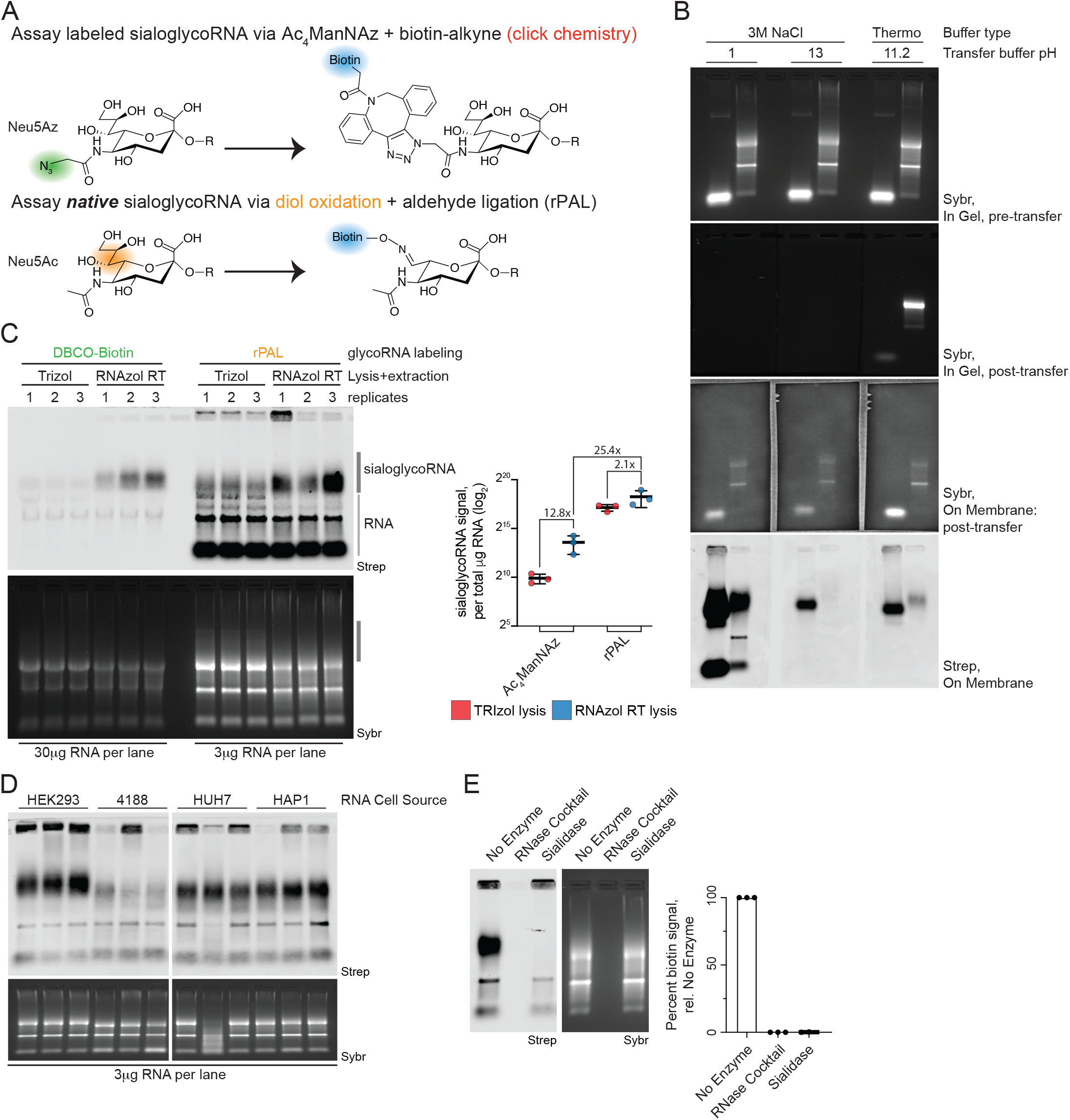
Selective oxidation and aldehyde labeling enables sensitive detection of native sialoglycoRNAs. A. Cartoon of sialic acid labeled strategies. Top: after uptake and conversion of Ac_4_ManNAz into Neu5Az, copper-free click chemistry can selectively label the Neu5Az with a biotin handle for detection. Bottom: native sialic acid diols can be oxidized and the resulting aldehyde ligated to a biotin reagent for detection (rPAL). B. RNA northern blot transfer optimization. Small and total RNA samples from HeLa cells labeled with rPAL were electrophoresed in a denaturing agarose gel and imaged with SybrGold (top row), after transferring with buffers of various pHs the gel was again imaged (second row) as well as the membrane where the RNA was transferred to (third row). Finally, sialoglycoRNAs were visualized with streptavidin-IR800 (Strep, bottom row). C. RNA blotting of HeLa total RNA labeled and detected with Ac_4_ManNAz paired with copper-free click of Dibenzocyclooctyne-PEG4-biotin (DBCO-biotin) or native sialic acids paired with rPAL labeling. Total RNA was extracted with either TRIzol or RNAzol RT to compare relative efficiencies of sialoglycoRNA detection. In gel detection of total RNA with SybrGold (Sybr, bottom) and on membrane detection of biotin (Streptavidin-IR800, top) is shown. The region labeled “sialoglycoRNA” was quantified and plotted (right). Each datapoint (biological triplicate) is displayed with the standard error of the mean (SEM). D. RNA blotting of HEK293, 4188, HUH7, and HAP1 total RNA. Detection of the resulting sialoglycoRNA signal was accomplished by rPAL labeling and quantified to the right. Sybr and Strep detection is as in C. E. RNA blotting of HeLa total RNA that was first treated with no enzyme, an RNase A and RNase T1 cocktail, or Sialidase. Detection of the resulting sialoglycoRNA signal was accomplished by rPAL labeling and quantified to the right. Sybr and Strep detection is as in C. Each datapoint (biological triplicate) is displayed with the SEM.

Ac_4_ManNAz enabled the glycoRNA discovery effort, but its limitations motivate the development of new tools. MCRs can only be taken up by metabolically active cells before their biosynthetic processing and incorporation into nascent biopolymers, like glycans. This limits MCR use in contexts where access to the organism or cells with a reporter is not possible. Glycan MCRs are also known to not fully saturate the sugar pool, resulting in the modification of only a portion of the total glycans^2^. Furthermore, glycan MCRs can be variably incorporated across glycoforms and cell types^2^. Additionally, the substitutions on MCRs (e.g. azide-group) can bias against their incorporation by certain glycosyltransferases^3^. Further characterization of glycoRNA species will necessitate the expansion of specific chemical tools that enable precise quantification, analysis of their dynamic cell display, and can be performed on the native glycoRNA substrate.

While methods exist to derivatize and label native glycoproteins and glycolipids, no such method exists for the modification of native glycoRNAs. We considered chemoenzymatic labeling, which requires charged sugar donors and specialized enzymes (Reviewed in^4^). Instead, we developed an RNA-optimized periodate oxidation and aldehyde labeling technique for the labeling of native glycoRNAs. We establish best practices for handling small RNA samples, develop a one-pot reaction for sialoglycoRNA labeling, and demonstrate its >25-fold increased sensitivity as compared to Ac_4_ManNAz. We further validate that periodate-labeled sialoglycoRNA behaves similarly to our previously reported Ac_4_ManNAz-labeling, establishing this method as an MCR-free alternative to study glycoRNAs. We anticipate our approach will accelerate the characterization of glycoRNAs structure and function across many biological contexts.

## Results

To enable detection of native sialoglycoRNAs, we considered the accessible functional groups within sialic acids. Existing methods applied to glycoproteins and glycolipids have leveraged the oxidation of diols vicinal to aldehydes and their subsequent ligation to amine-containing reagents or supports. Paired with aminooxy containing molecules, this reaction forms a stable oxime bond without requiring further derivatization. While this approach is reasonably selective for sialic acid-containing glycans, the presence of a 2’, 3’ vicinal diol at the 3’ terminal ribose of RNA poses a challenge to its application to glycoRNA. While the 7’, 8’ diols in sialic acid are quite reactive—and thus can be derivatized at physiological pH with short reaction times^5^—complete oxidation of ribose diols in RNA requires more acidic buffers and longer reaction times^6^. With this insight into the differential reactivity of sialic acid diols and ribose diols, we reasoned that in a mixture of RNA nucleotides and sialic acid sugars, mild oxidation conditions could achieve selective sialic acid diol labeling (**Figure 1A**).

Over the course of this work, we noticed that precise handling, specific reagents, and optimized procedures dramatically improved reproducibility across our experiments. These features are detailed in the Methods section, but major points are noted here. Our standard cell lysis used TRIzol, however other reagents like RNAzol RT have been developed to use less phenol and no chloroform in the denaturing and RNA extraction processes^7^. RNAzol RT extraction of HeLa cells resulted in similar amounts of total RNA isolated per cell compared to TRIzol (**Figure S1A**). Column clean ups are convenient for rapid processing and are reported to have high RNA recovery. We found that under recommended conditions, HeLa cell total RNA recovery was ∼92% (**Figure S1B**) while small RNA recovery was < 60% (**Figure S1C**). We hypothesized this was an issue with the precipitation strength of the prescribed binding reaction and tested various alcohol types and ratios, finding that both ethanol and isopropanol at adjusted ratios could facilitate > 90% recovery of small RNAs using Zymo columns (**Figure S1C**). With a more reproducible means to isolate and re-purify RNA across chemical reactions, we proceeded to develop a method to label native sialoglycoRNAs.

To maintain a simple procedure, we set out to develop a reaction scheme that would both oxidize diols and ligate newly generated aldehydes (periodate oxidation and aldehyde labeling, PAL) to a labeling reagent in the same reaction without purification. Conditions including pH, salt concentration, salt type, and temperature were screened using RNAzol RT extracted HeLa cell total RNA (**Figure S1D, S1E, S1F**). An RNA optimized PAL protocol (rPAL, **Methods**) includes a pre-blocking step with a free aldehyde reagent which reduces background signal in the well (**Figure S1G**) and takes in total 2.5 hours. During the development of this protocol we re-assessed the need for protease digestion, RNA was extracted without the assistance of proteinase K or StcE during the preparation and only after RNA was isolated, did we then add enzyme to determine co-purification of background proteins and mucins. As we saw previously, proteinase K is important to fully remove proteins from isolated RNA, however while Ac_4_ManNAz signal was not sensitive to mucinase (StcE) digestion^1^ the PAL signal was (**Figure S1H**). Additionally, we noticed that our commercially sourced RNA transfer buffer produced incomplete RNA transfers even under the prescribed conditions (**Figure 1B** and **S2A**). To establish the impact of this, we first screened the pH of the buffer on the transfer properties of Ac_4_ManNAz-labeled RNA. We found that pH < 2 or > 12 led to complete RNA transfer and increased the amount of glycoRNA signal deposited on the membrane (**Figure S2A**). This increase was apparent but mild and we also noticed additional background bands transferred from Ac_4_ManNAz-labeled RNA at low pH (**Figure S2A**). We next tested the transfer buffers to verify complete deposition of the rPAL signal onto nitrocellulose membranes. Unlike the ManNAz-labeled signal, which was similarly enhanced at high and low pH, the rPAL signal was slightly weaker using the basic pH buffer conditions while its transfer was dramatically enhanced at low pH (**Figure 1B**). Further testing defined the optimal pH, salt, and time parameters (**Figure S2B and S2C**), resulting in an optimized method and enhanced transfer conditions as well as allowing us now to directly compare Ac_4_ManNAz signal to rPAL signal.

Extracting RNA from Ac_4_ManNAz labeled HeLa cells with TRIzol and either performing copper-free click^1^ or rPAL, we found that rPAL generates approximately 150x the amount of signal (**Figure 1C**, lanes 1-3 vs 7-9). We repeated this comparison from Ac_4_ManNAz-labeled cells and while there was not a significant difference in total RNA extracted with TRIzol vs RNAzol RT (**Figure S1A**), we saw a 12.8x and 2.1x fold gain in glycoRNA signal with Ac_4_ManNAz and rPAL respectively (**Figure 1C**). Integrating both the rPAL and RNAzol RT RNA extraction, we can achieve at least 25x fold increased signal recovery per mass of RNA compared to an updated Ac_4_ManNAz-strategy that uses RNAzol RT extractions (**Figure 1C**). While rPAL improves sensitivity of apparent high molecular weight (MW) glycoRNA species, it also induces background labeling; most notably the 18S rRNA and the small RNA pool (**Figure 1C** and elsewhere). This is expected given that the 3’ ends of RNAs should (mostly) contain a 2’-3’ vicinal diol, which can also undergo periodate-based oxidation^6^. To demonstrate the flexibility of rPAL, we labeled total RNA stocks which were collected over 3 years ago from cells treated with Ac_4_ManNAz for 24 hours prior to collection. rPAL readily detects sialoglycoRNAs from the four archived RNA stocks sourced from HEK293, 4188, HUH7, and HAP1 cells (**Figure 1D**). We can more directly interpret the levels of rPAL signal across each cell type as changes in the sialylation of glycoRNA glycans, unlike with Ac_4_ManNAz-labeling where the metabolism and incorporation of Ac_4_ManNAz must also be considered. Finally, to establish the high MW signal is indeed sialoglycoRNA, we evaluated its sensitivity to enzymatic digestion. Incubation of purified RNA with RNase or sialidase and subsequent rPAL-labeling results in total loss of all rPAL signal after RNase but selective and complete loss of the high MW signal after sialidase treatment (**Figure 1E**), consistent with this rPAL-based method labeling sialic acids in sialoglycoRNAs.

In our initial report of sialoglycoRNAs as detected by Ac_4_ManNAz, we described several key features including that small RNAs were the template, that they fractionated with cellular membranes, and that they are accessible to the surface of living cells^1^. To verify that the high MW signal is indeed small glycoRNAs, we used column-based separation of large and small RNAs and found the high MW rPAL signal, as well as the signal overlapping the small RNA pool, is enriched with the small RNAs but not the long RNAs (**Figure 2A**). Assessment of the subcellular localization demonstrated that the high MW signal co-purifies with crude cellular membrane extracts while it is depleted in the cytosolic fraction (**Figure 2B and S3A**). Additionally, and in line with our reported topology of Ac_4_ManNAz-labeled sialoglycoRNAs, rPAL-labeled high MW signal is sensitive to live cell sialidase treatment. By treating live HeLa cells with vibrio cholerae (VC) sialidase, extracting RNA, and performing rPAL labeling, we could detect robust loss of sialoglycoRNA signal in as little as 15 minutes after addition of the VC-sialidase (**Figure 2C**).

**Figure 2.**
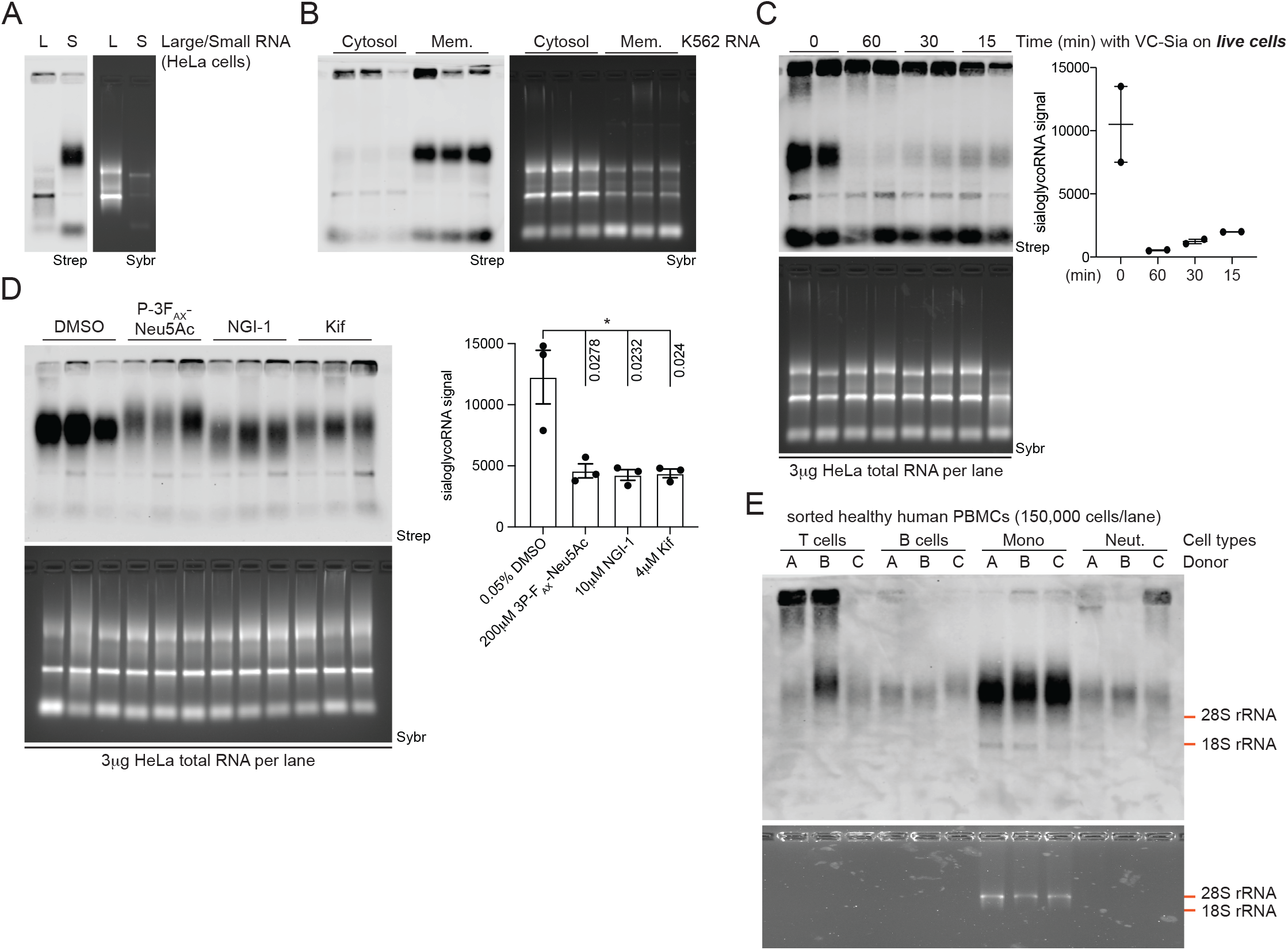
rPAL detects small, cell surface-exposed sialoglycoRNAs sensitive to inhibitors of N-glycosylation. A. RNA blotting of HeLa RNA that after isolation was subjected to a size fractionation, separating out large (L, > ∼200 nts) from small (S, < ∼200 nts) RNAs. Detection of the resulting sialoglycoRNA signal was accomplished by rPAL labeling, in gel detection of total RNA with SybrGold (Sybr, bottom) and on membrane detection of biotin (Streptavidin-IR800, top) is shown. B. RNA blotting of RNA that was isolated from K562 cells fractionated for their crude membranes. Sybr and Strep detection is as in A. C. RNA blotting of total RNA from HeLa cells that prior to RNAzol RT lysis were exposed to VC-Sialidase treatment in the culture media for the indicated times. After treatment, cells were washed twice with 1x PBS, and then RNA collected and labeled with rPAL. Sybr and Strep detection is as in A. SialoglycoRNA signal was quantified and plotted (right). Each datapoint (biological duplicate) is displayed with the SEM. D. RNA blotting of total RNA from HeLa cells treated with various chemical inhibitors. Inhibitors were added to cells in complete media for 24 hours: 0.05% DMSO, 200μM P-3_FAX_-Neu5Ac, 10μM NGI-1, or 4μM Kifensuesine, after which RNA was collected and processed for rPAL labeling. Sybr and Strep detection is as in A. SialoglycoRNA signal was accomplished by rPAL labeling and quantified to the right; each datapoint (biological triplicate) is displayed with the SEM and statistical analysis was performed using an unpaired t-test. E. RNA blotting of total RNA from four sorted populations of human PBMCs including T cells, B cells, Monocytes (Mono) and Neutrophils (Neut.). RNA from approximately 150,000 cells were used for each lane across the four populations.

Labeling sialoglycoRNAs with Ac_4_ManNAz resulted in a signal that was sensitive to inhibition of enzymes responsible for N-glycosylation. We next determined how the rPAL-signal from HeLa cells responded to inhibition of sialic acid biosynthesis with P-3F_AX_-Neu5Ac^8^, the oligosaccharyltransferase with NGI-1^9^, or ɑ-mannosidase-I with Kifunensine^10^. In all three cases we saw loss of the rPAL-signal (**Figure 2D**), consistent with our previous results detecting sialoglycoRNAs with Ac_4_ManNAz. P-3F_AX_-Neu5Ac and Kifunensine treatment caused a larger and milder shift higher in the apparent MW of the sialoglycoRNAs (**Figure 2D**), respectively, which is in line with the effects seen with Ac_4_ManNAz. NGI-1 resulted in an apparent lower MW smearing of the signal (**Figure 2D**), which was not something we had previously seen with Ac_4_ManNAz.

To demonstrate the utility of the significantly increased sensitivity of rPAL and the ability to access samples where MCR-labeling would be challenging, we examined the sialoglycoRNAs from human peripheral blood mononuclear cells (PBMCs). We stained healthy human PBMCs with antibodies targeting CD19 (B cells), CD3 (T cells), CD14 (Monocytes), and CD16 (Neutrophils) and performed cell sorting (**Figure S3B**). We sorted 150,000 cells of each cell type per tube per donor (three total donors) after which RNA was extracted (total RNA amounts in the 100’s of nanograms per reaction) and processed with the rPAL labeling method. Across all four cell types sorted we can detect sialoglycoRNAs (**Figure 2E**). All donors generated sialoglycoRNA signal from the four cell types analyzed, with the sorted monocytes both yielding the most total RNA as well as the strongest sialoglycoRNA signal (**Figure 2E**). Some samples from each of the cell types also develop ultrahigh MW signal (**Figure 2E**), which is evident but less common in material isolated from cultured lines with higher cell numbers (e.g. Figure 2C, lanes 1, 3, and 4). We note that minor MW change could be seen between donors of T cells (donor 3) and B cells (donor 2) (**Figure 2E**).

## Discussion

Here we describe the development of “rPAL”, which is a simple, one pot, rapid, and sensitive means to label native sialoglycoRNAs from isolated RNA samples. Leveraging rPAL enables detection of sialoglycoRNAs without the need to incorporate metabolic chemical reporters, opening a greater set of biological sources for investigation of sialoglycoRNA biology. Changes in sialoglycoRNA signal can now be more directly linked to the biosynthetic flux of sialyltransferases rather than possible differences from cell-type specific variations in metabolic reporter uptake and intracellular usage of Ac_4_ManNAz^11^. Additionally, coupled with the > 25x increase in sensitivity over Ac_4_ManNAz-based detection, rPAL allows for analysis of primary human samples for which Ac_4_ManNAz labeling is not easily accomplished and where the cell number can be limiting. Our investigation into major immune effector cells isolated from primary human PBMCs and sorted with FACS demonstrated detection of sialoglycoRNAs produced in T cells, B cells, Monocytes, and Neutrophils. This was accomplished with only 150,000 cells and enabled by the significantly increased sensitivity of rPAL coupled to the improved RNA processing and RNA blotting strategies discussed below. This FACS-based workflow will allow for investigation of other precious samples such as material collected from patients to understand how sialoglycoRNAs change in disease. Future work that expands the set of monosaccharides (beyond sialic acid) that can be labeled and investigated in the context of glycoRNA biology will further reveal features of how glycoRNAs operate. Additionally, we see opportunities to directly visualize glycoRNAs with imaging techniques as an exciting future direction.

Over the course of this work, a series of technical advances were also achieved that should be broadly applicable to other aspects of RNA biology. First, silica-based columns offer a convenient and rapid means to clean nucleic acids up between reactions; however, we found that while they are highly efficient for large RNAs, short RNAs are poorly recovered under standard conditions. By exploring various binding conditions, we have now optimized conditions that are nearly quantitative for small RNA capture and elution (**Figure S1A-C**), allowing serial reaction processing without major losses in material. This has improved sample-to-sample variability as well as enabled lower starting material procedures. Second, detection of biomolecules with probes or other protein affinity tools fail when the physical transfer from 2D gel separation onto membranes is inefficient. We found that standard methods and buffers to transfer RNA out of denaturing agarose gels were not effective, particularly for rPAL-labeled glycoRNAs. By screening various salt and pH conditions we found a highly efficient process to fully transfer all of the total RNA as well as significantly more sialoglycoRNAs onto the nitrocellulose membrane compared with commercial buffer stocks. Importantly, improved overall transfer of Ac_4_ManNAz-signal resulted in background bands appearing (**Figure S2A**, left most column) which mirrors some of the background labeling of total RNA we find with rPAL labeling (**Figure 1C**, “RNA” label). Thus, while neither rPAL or Ac_4_ManNAz is totally background free, the unusual migration of sialoglycoRNAs in the agarose gels enables facile interpretation. Finally, while screening for efficient transfer of Ac_4_ManNAz-vs rPAL-signal, we noticed that Ac_4_ManNAz-signal was enhanced at both high and low pH, while rPAL was transferred poorly at high pH and very efficiently at low pH (**Figure S2**). This is another difference between Ac_4_ManNAz and rPAL, and suggests some compositional differences in the bulk glycans that are being detected with each method.

We used Ac_4_ManNAz-labeled sialoglycoRNAs as a benchmark, and based on our data, the rPAL-labeled sialoglycoRNAs behave similarly but not identically. rPAL signal fractionated with small RNAs, was completely sialidase sensitive, was reduced in accumulation in cells after inhibiting N-glycosylation biosynthetic enzymes, and importantly rPAL labeled sialoglycoRNAs are highly sensitive to live cell treatment with sialidase, confirming cell surface presentation. However, there are some new features we uncovered using this labeling method. First and most obviously, rPAL labels significantly more sialoglycoRNAs per mass of total RNA from cells, suggesting that Ac_4_ManNAz was relatively inefficient from a fractional incorporation standpoint. The ability to label fractionally more of the sialoglycoRNAs in the cell opened the possibility that we may be detecting novel sialoglycoRNAs in addition to the forms found initially with Ac_4_ManNAz. For example, while we noticed clear and significant loss of rPAL signal after NGI-1 and Kif treatment of HeLa cells, the effects were less than what we had previously seen only with Ac_4_ManNAz-labeling^1^. Additionally, we previously tested Ac_4_ManNAz-signal for sensitivity to mucinase digestion and saw no effect, while unprocessed RNA labeled with rPAL is partially sensitive to mucinase treatment (**Figure S1H**). These differences suggest that rPAL labeling sees deeper into the set of sialoglycoRNAs that are generated in each cell, and highlights a possibility that there is an expanded set of glycoforms modifying RNA or RNAs being modified by sialoglycans. The facile nature of the rPAL method should enable new investigations of glycoRNAs with lower effort and in a broader set of biological contexts.

## Methods

### Cell culture, chemical inhibitor, and metabolic chemical reporters

All cells were grown at 37°C and 5% CO_2_. HeLa cells were cultured in DMEM media supplemented with 10% fetal bovine serum (FBS) and 1% penicillin/streptomycin (P/S). RNA from other cell sources were obtained from^1^ and re-purified as per the descriptions below. Stocks of N-azidoacetylmannosamine-tetraacylated (Ac_4_ManNAz, Click Chemistry Tools) were made to 500 mM in sterile dimethyl sulfoxide (DMSO). For cell treatments, Ac_4_ManNAz was used at a final concentration of 100 μM. Working stocks of glycan-biosynthesis inhibitors were all made in DMSO at the following concentrations and stored at -80°C: 5 mM NGI-1 (Sigma), 10 mM Kifunensine (Kif, Sigma), 50 mM P-3F_AX_-Neu5Ac (Tocris). All compounds were used on cells for 24 hours.

### RNA extraction and enzymatic cleanups

TRIzol extractions were performed as previously described in detail^1^. For RNAzol RT (Molecular Research Center, Inc.) extractions, the manufacturer’s protocol was followed with the following details. First, RNAzol RT was added to lyse and denature cells or tissues, and denaturing was further encouraged by placing the samples at 50°C and shaking for 5 min. To phase separate the RNA, 0.4X volumes of water was added, vortexed, let to stand for 5 minutes at 25°C and lastly spun at 12,000x g at 4°C for 15 min. The aqueous phase was transferred to clean tubes and 1.1X volumes of isopropanol was added. The RNA is then purified over a Zymo column (Zymo Research). We found that preconditioning Zymo columns with water before binding nucleic acids produces more consistent results. For all column cleanups, we followed the following protocol. First, 350μL of pure water was added to each column and spun at 10,000x g for 30 seconds, and the flowthrough was discarded. Next, precipitated RNA from the RNAzol RT extraction (or binding buffer precipitated RNA, below) is added to the columns, spun at 10,000x g for 10-20 seconds, and the flowthrough is discarded. This step is repeated until all the precipitated RNA is passed over the column once. Next, the column is washed three times total: once using 400μL RNA Prep Buffer (3M GuHCl in 80% EtOH), twice with 400μL 80% ethanol. The first two spins are at 10,000x g for 20 seconds, the last for 30 sec. The RNA is then treated with Proteinase K (Ambion) on the column. Proteinase K is diluted 1:19 in water and added directly to the column matrix (Zymo-I = 20μL, Zymo-II = 50μL, Zymo-IIICG = 60μL, all from Zymo Research), and then allowed to incubate on the column at 37°C for 45 min. The column top is sealed with either a cap or parafilm to avoid evaporation. After the digestion, the columns are brought to room temperature for 5 min; lowering the temperature is important before proceeding. Next, eluted RNA is spun out into fresh tubes and a second elution with water is performed (Zymo-I = 30μL, Zymo-II = 50μL, Zymo-IIICG = 60μL). To the eluate, 1.5μg of the mucinase StcE (Sigma-Aldrich) is added for every 50μL of RNA, and placed at 37°C for 30 minutes to digest. The RNA is then cleaned up again using a Zymo column. Here, 2X RNA Binding Buffer (Zymo Research) was added and vortexed for 10 seconds, and then 2X (samples + buffer) of 100% ethanol was added and vortexed for 10 sec. An example would be 50μL of RNA, 100μL of RNA Binding Buffer, and 300μL of 100% ethanol. This is then bound to the column, cleaned up as described above, and eluted twice with water (Zymo-I = 25μL, Zymo-II = 50μL, Zymo-IIICG = 60μL). The final enzymatically digested RNA is quantified using a Nanodrop.

Binding conditions to efficiently precipitate small RNAs as highlighted in **Figure S1C** were optimized by varying the amount of ethanol added post RNA Binding Buffer mixing with RNA. Isopropanol was also exchanged for the ethanol at this step to assess its ability to facilitate small RNA capture on the Zymo columns.

After total RNA extraction, the RNA can be further processed in order to fractionate small (17-200 nts) and large RNA (>200 nts) using Zymo columns. First, an adjusted RNA binding buffer is made by mixing equal volumes of RNA binding buffer and 100% ethanol. Two volumes of the adjusted buffer are added to the total RNA and are vortexed thoroughly to mix. The sample is then bound to the column as described above, but the flow through - which contains the small RNA - is saved. One volume of 100% ethanol is added to the small RNA and vortexed to mix. The small RNA is then bound to new columns. Both the large and small RNA (bound to their respective columns) are then cleaned up using 2X 400μL 80% ethanol, the second clean being centrifuged for 30 seconds. The RNA is then eluted using 2X 50μL of water.

### In vitro and On Cell enzymatic digestions

To digest RNA, the following was used: 2μL of RNase cocktail (0.5U/mL RNaseA and 20U/mL RNase T1, Thermo Fisher Scientific) with 20 mM Tris-HCl (pH 8.0), 100 mM KCl and 0.1 mM MgCl_2_. To digest sialic acid: 1.5μL of a2-3,6,8 Neuraminidase (50U/mL, New England Biolabs, NEB) with 1x GlycoBuffer 1 (NEB). Reactions were performed on 3μg total RNA from indicated cell sources for 60 minutes at 37°C. For live cell treatments, VC-Sia was expressed and purified as previously described (Gray et al., 2019) and added to cells at 150 nM final concentration in complete growth media for between 15 and 60 minutes at 37°C.

### Isolation of crude cellular membranes

Crude membranes were isolated using the Plasma Membrane Protein Extraction Kit (ab65400, Abcam): cultured cells first had growth media removed and cells were then washed twice with ice-cold 1x Phosphate Buffered Saline (PBS). In the second PBS wash, cells were scraped off the plate and spun down at 400x g for 4 minutes at 4°C or suspension cells were directly pelleted. Cell pellets were resuspended in 1mL of Homogenization Buffer Mix per 10,000,000 cells. Cell suspension was Dounce Homogenized on ice for 40-70 strokes, care was taken to stop douncing when the processing resulted in approximately 60% free nuclei so as to not generate excess lysis of nuclei. Homogenate was then spun at 700x g for 10 minutes at 4°C. This pellet contained the nuclear fraction and supernatants were transferred to new tubes and spun again at 10,000x g for 30 minutes at 4°C. The pellets generated from this spin were crude membranes and the supernatant was soluble cytosol. RNA extraction was performed as above and labeling as described below.

### Periodate oxidation and aldehyde ligation for glycoRNA labeling

Starting with a maximum of 3ug of lyophilized, enzymatically treated RNA, the first step of the rPAL labeling protocol is to block any aldehyde reactive species. To make the blocking buffer, 1μL 16 mM mPEG3-Ald (BP-23750, BroadPharm), 15μL 1 M MgSO_2_ and 12μL 1 M NH_4_OAc pH5 (with HCl) are mixed together (final buffer composition: 570 μM mPEG3-Ald + 500 mM MgSO_4_ + 450 mM NH_4_OAc pH5). 28μL of the blocking buffer is added to the lyophilized RNA, mixed completely by vortexing, and then incubated for 45 minutes at 35°C to block. The samples are briefly allowed to cool to room temperature (2-3 min), then working quickly, 1μL 30 mM aldehyde reactive probe (ARP, aminooxy) biotin (22582, Cayman Chemicals, stock made in water) is added first, then 2μL of 7.5 mM NaIO_4_ (periodate, stock made in water) is added. The periodate is allowed to perform oxidation for exactly 10 minutes at room temperature in the dark. The periodate is then quenched by adding 3μL of 22 mM sodium sulfite (stock made in water). The quenching reaction is allowed to proceed for 5 minutes at 25°C. Both the sodium periodate and sodium sulfite stocks were made fresh within 20 minutes of use. Next, the reactions are moved back to the 35°C heat block, and the ligation reaction is allowed to occur for 90 min. The reaction is then cleaned up using a Zymo-I column. 19μL of water is added in order to bring the reaction volume to 50μL, and the Zymo protocol is followed as per the above details. If samples will be analyzed on an agarose gel for glycoRNA visualization, the RNA is then eluted from the column using 2X 6.2μL water (final volume approximately 12μL).

### glycoRNA blotting

In order to visualize the periodate labeled RNA, it is run on a denaturing agarose gel, transferred to a nitrocellulose (NC) membrane, and stained with streptavidin in a manner similar to Flynn et al. 2021^1^ with some modifications. After elution from the column as described above, the RNA is combined with 12μL of Gel Loading Buffer II (GLBII, 95% formamide, 18 mM EDTA, 0.025% SDS) with a final concentration of 2X SybrGold (ThermoFisher Scientific) and denatured at 55°C for 10 minutes. It is important to not use GLBII with dyes. Immediately after this incubation, the RNA is placed on ice for at least 2 minutes. The samples are then loaded into a 1% agarose, 0.75% formaldehyde, 1.5x MOPS Buffer (Lonza) denaturing gel. Precise and consistent pouring of these gels is critical to ensure a similar thickness of the gel for accurate transfer conditions; we aim for approximately 1 cm thick of solidified gel. RNA is electrophoresed in 1x MOPS at 115V for between 34 or 45 min, depending on the length of the gel. Subsequently, the RNA is visualized on a UV gel imager, and excess gel is cut away; leaving ∼0.75 cm of gel around the outer edges of sample lanes will improve transfer accuracy. The RNA is transferred as previously described^1^, however various buffer conditions and times were screened to determine the optimal method (**Figure S2**). Finally, we determined that a 3M NaCl solution at pH 1, achieved with HCl, yields the most consistent and efficient transfer of material to the NC membrane. Transfer occurs for 90 minutes at 25°C. Post transfer, the membrane is rinsed in 1x PBS and dried on Whatman Paper (GE Healthcare). Dried membranes are rehydrated in Intercept Protein-Free Blocking Buffer, TBS (Li-Cor Biosciences), for 30 minutes at 25°C. After the blocking, the membranes are stained using Streptavidin-IR800 (Li-Cor Biosciences) diluted 1:5,000 in Intercept blocking buffer for 30 minutes at 25°C. Excess Streptavidin-IR800 was washed from the membranes using three washes with 0.1% Tween-20 (Sigma) in 1x PBS for 3 minutes each at 25°C. The membranes were then briefly rinsed with PBS to remove the Tween-20 before scanning. Membranes were scanned on a Li-Cor Odyssey CLx scanner (Li-Cor Biosciences).

### Peripheral blood mononuclear cell staining and flow cytometry sorting

Human Peripheral Blood Mononuclear Cells (PBMCs) were purchased from StemCell Technology. Samples from three individual donors were thawed at 37°C for 3 minutes and then diluted into 5 mL of FACS buffer (0.5% BSA in 1x PBS). Freezing media was removed by spinning the cells for 4 minutes at 400g at 4°C. 10 million cells were resuspended in 1 mL of FACS buffer and 50 μL of Human TruStain FcX Fc Receptor Blocking Solution (BioLegend) was added and left to bind the cells on ice for 15 minutes. The blocked PMBCs were stained with anti-CD19 clone HIB19 conjugated to PE-Cy7 (BioLegend), anti-CD3 clone OKT3 conjugated to FITC (BioLegend), anti-CD14 clone HCD14 conjugated to APC-Cy7 (BioLegend), and anti-CD16 clone 3G8 conjugated to PerCP-Cy5.5 (BioLegend) for 30 minutes on ice. Cells were pelleted by spinning for 4 minutes at 400g at 4°C. The supernatant was discarded and the cells were resuspended in 1 mL of FACS buffer supplemented with 4’,6-Diamidino-2-phenylindole (DAPI) at 1 μg/mL final concentration as a live/dead stain. Stained cells were sorted into four major subpopulations using a Sony MA900 sorter with a 100 μM sorting chip (Sony). Cells were collected into FACS buffer and spun for 4 minutes at 400g at 4°C. Pellets were lysed in RNAzol RT and rPAL labeling was performed as described above.

## Figure Legends

**Figure S1.**
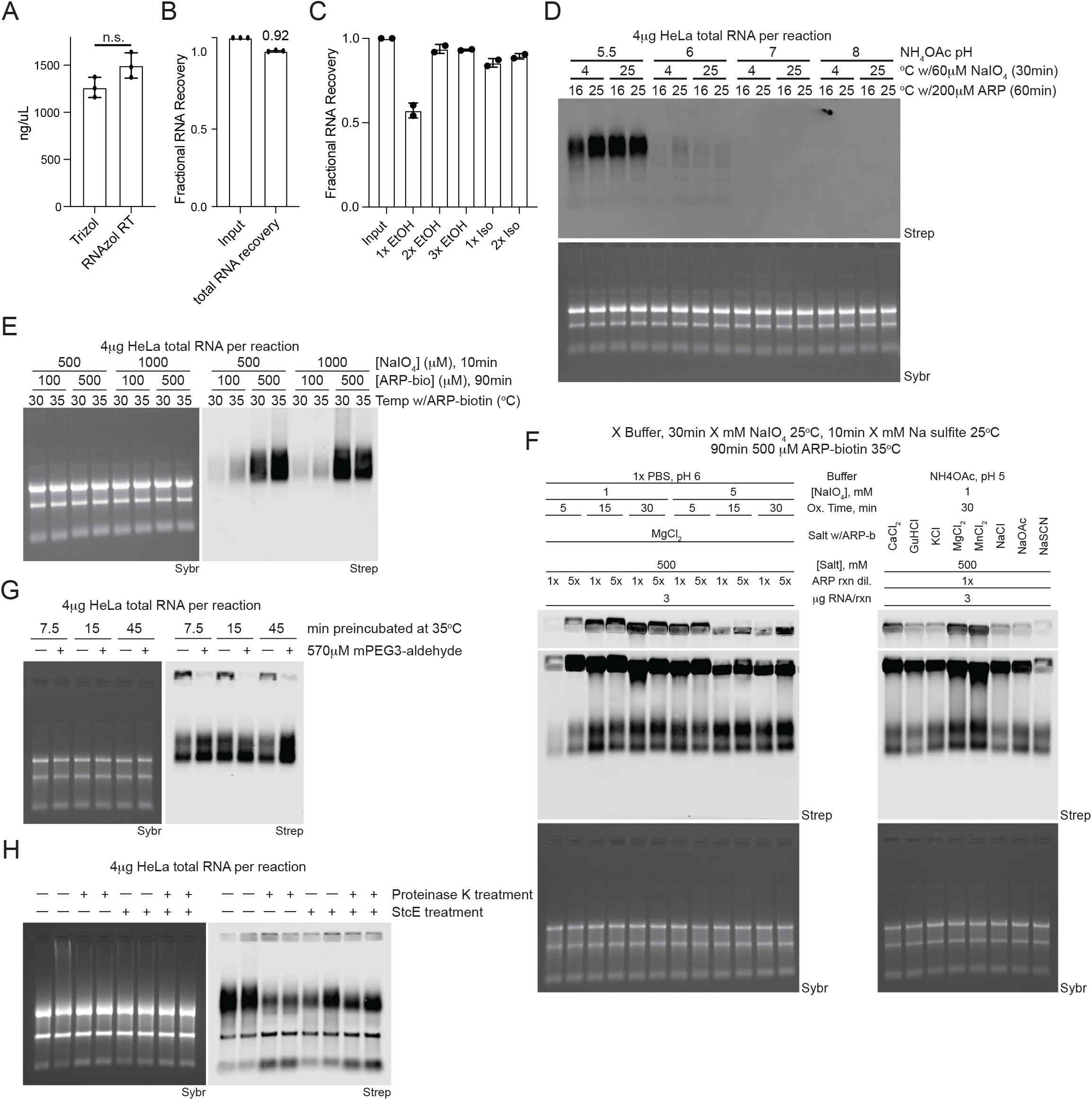
Development of rPAL labeling strategy. A. Nanodrop analysis of total RNA extracted from HeLa cells. Cells were lysed with either Trizol or RNAzol RT and processed as per the manufacturer’s recommendations with RNA cleanups performed using Zymo columns (**Methods**). Each datapoint (biological triplicate) is displayed with the SEM. B. Nanodrop analysis of total RNA extracted from HeLa cells isolated with RNAzol RT. After isolation, samples were quantified and then subjected to a Zymo column clean up after which each sample was quantified again. Each datapoint (biological triplicate) is displayed with the SEM. C. Nanodrop analysis of small RNA extracted from HeLa cells isolated with RNAzol RT. After isolation, samples were quantified and then subjected to a Zymo column clean up. During the precipitation step, before binding to the Zymo columns, indicated ratios of ethanol or isopropanol were added. After Zymo column cleanup, each sample was quantified again. Each datapoint (biological duplicate) is displayed with the SEM. D. RNA blotting of total RNA from HeLa cells. Total RNA was subjected to various reaction conditions including NH_4_OAc buffer pH changes, the temperature at which diol oxidation occurred, and the temperature at which the aldehyde ligation occurred. In gel detection of total RNA with SybrGold (Sybr, bottom) and on membrane detection of biotin (Streptavidin-IR800, top) is shown. E. Blotting as in D with variations in the concentration of NaIO_4_ used for oxidation and ARP-biotin used for aldehyde ligation as well as the temperature at which the aldehyde ligation occurred at. Sybr and Strep detection is as in D. F. Blotting as in D with evaluation of buffer conditions, NaIO_4_ concentration, ARP-biotin concentrations, and oxidation times for generating specific rPAL signal. Sybr and Strep detection is as in D. G. Blotting as in D with evaluation of performing a blocking step before the NaIO_4_ oxidation with mPEG3-aldehyde for the indicated times at 35°C. Sybr and Strep detection is as in D. H. Blotting as in D with evaluation of enzymatic digestions of the RNA samples after extraction using RNAzol RT. RNA was either not digested or subjected to digestion with Proteinase K (45 minutes at 37°C), StcE (35 minutes at 37°C), or both enzymes (sequentially). Sybr and Strep detection is as in D.

**Figure S2.**
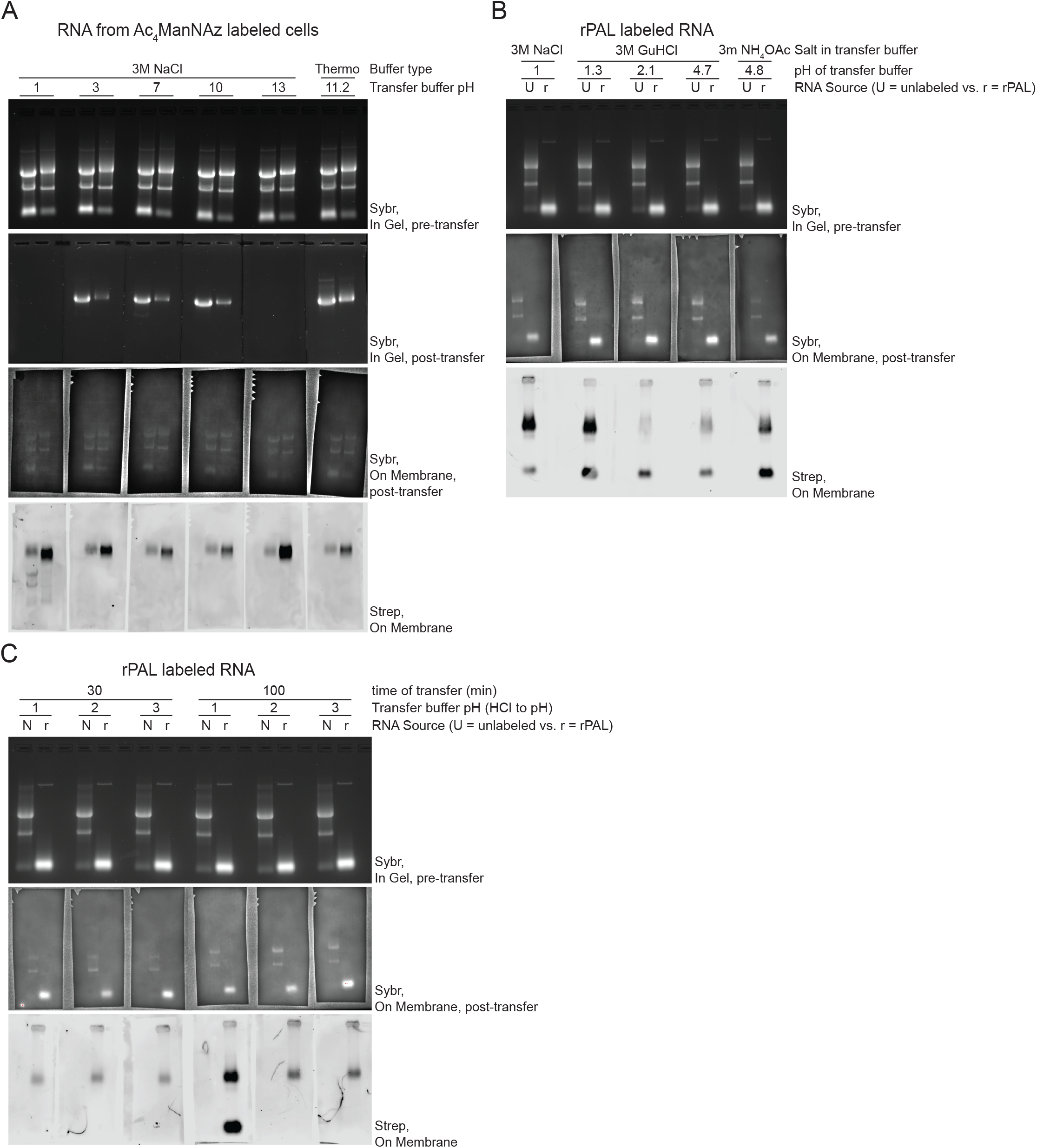
Optimization of RNA Northern blotting transfer conditions. A. RNA northern blot transfer optimization. Small and total RNA samples from HeLa cells labeled with Ac_4_ManNAz and visualized with copper-free click of DBCO-biotin were electrophoresed in a denaturing agarose gel and imaged with SybrGold (top row), after transferring with buffers of various pHs the gel was again imaged (second row) as well as the membrane where the RNA was transferred to (third row). Finally, sialoglycoRNAs were visualized with streptavidin-IR800 (Strep, bottom row). B. RNA northern blot transfer optimization as in A, screening buffer salt composition as well as pH. Here the detection of sialoglycoRNAs was accomplished with rPAL. C. RNA northern blot transfer optimization as in B, assessing the time and pH dependency of transfer efficiency.

**Figure S3.**
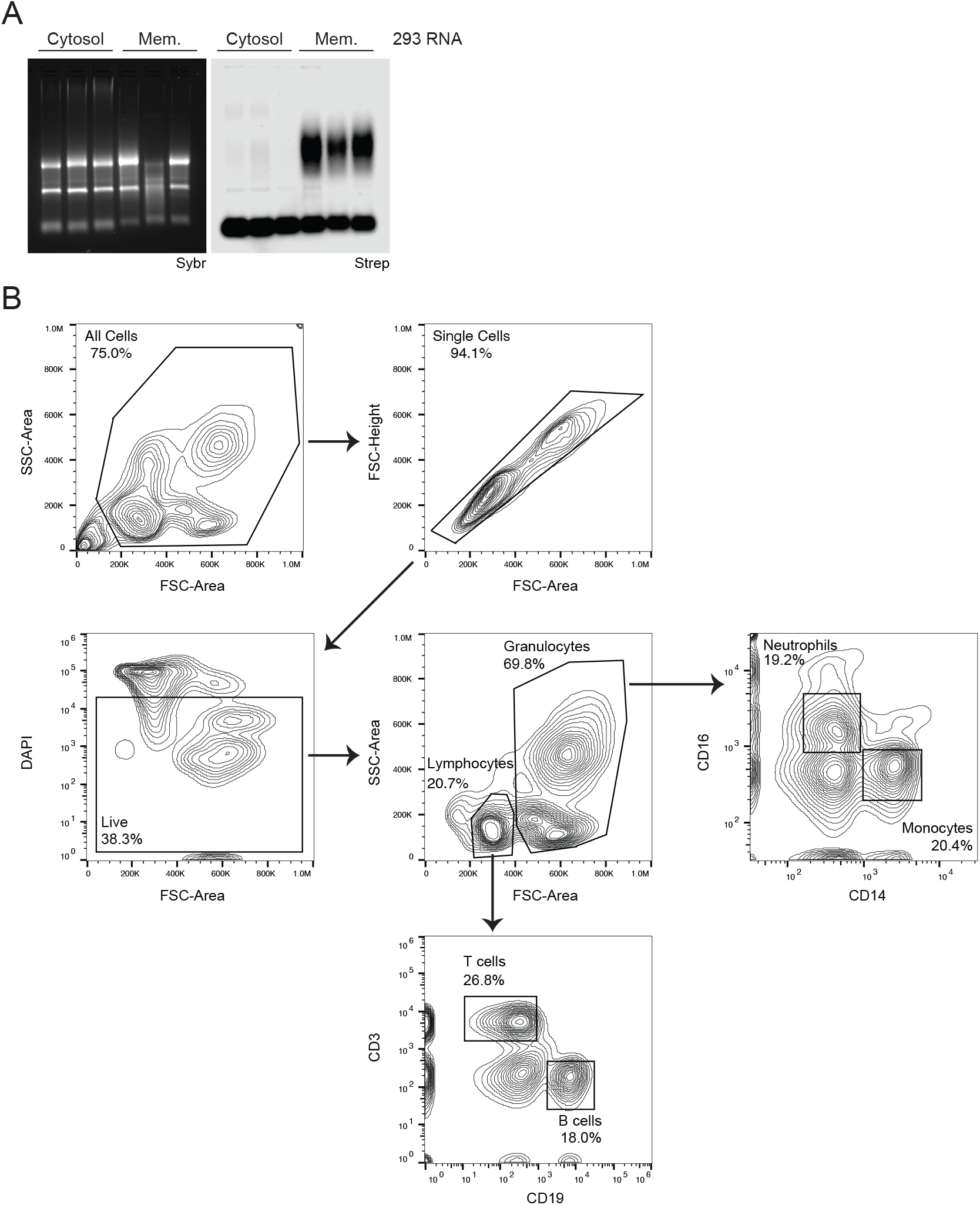
Subcellular localization of rPAL signal and flow cytometry analysis for sorting of human PBMCs. A. RNA blotting of RNA that was isolated from HEK293 cells fractionated for their crude membranes. The soluble cytosol is a fraction from this procedure that serves as a non-membrane enriched control. B. Scatter plots of one of three human peripheral blood mononuclear cell (PBMC) sorting experiments. Each plot represents serial gates drawing to isolate live single cells and further fractionation T cells, B cells, Monocytes, and Neutrophils. These four populations were then sorted into FACS Buffer for later RNA extraction and sialoglycoRNA analysis. Arrows denote the gating scheme used to isolate the four final populations of cells.

## Acknowledgments

We thank Kayvon Pedram, Carolyn R. Bertozzi, and Robert C. Spitale for helpful comments and discussions. We also thank Melissa Gray for the expression and purification of VC-Sialidase. This work was supported by grants from Burroughs Wellcome Fund Career Award for Medical Scientists (R.A.F.), The Sontag Foundation Distinguished Scientist Award (R.A.F.), The Rita Allen Foundation (R.A.F.), a private donation administered by the National Philanthropic Trust (R.A.F.), and the Herchel Smith-Harvard Undergraduate Science Research Program (R.M.C.).

## Author Contributions

R.A.F. conceived and supervised the project. H.H., P.C., C.G.L., R.M.C., and R.A.F. performed RNA labeling optimization experiments and cell-based assays. B.M.G. performed PBMC staining and FACS sorting. R.A.F. wrote the manuscript. All authors discussed the results and revised the manuscript.

